# Intra-colony channel morphology in *Escherichia coli* biofilms is governed by nutrient availability and substrate stiffness

**DOI:** 10.1101/2021.12.17.473116

**Authors:** Beatrice Bottura, Liam M. Rooney, Paul A. Hoskisson, Gail McConnell

**Affiliations:** Department of Physics, SUPA, University of Strathclyde, G4 0NG, Glasgow, UK; Strathclyde Institute of Pharmacy and Biomedical Sciences, University of Strathclyde, G4 0RE, Glasgow, UK

## Abstract

Nutrient-transporting channels have been recently discovered in mature *Escherichia coli* biofilms, however the relationship between intra-colony channel structure and the surrounding environmental conditions is poorly understood. Using a combination of fluorescence mesoscopy and a purpose-designed open-source quantitative image analysis pipeline, we show that growth substrate composition and nutrient availability have a profound effect on the morphology of intra-colony channels in mature *E. coli* biofilms. Under all nutrient conditions, intra-colony channel width was observed to increase non-linearly with radial distance from the centre of the biofilm. Notably, the channels were around 25% wider at the centre of glucose-limited biofilms compared to ammonium-limited biofilms. Channel density also differed in colonies grown on rich and minimal media, with the former creating a network of tightly packed channels and the latter leading to well-separated, wider channels with defined edges. Our approach paves the way for measurement of internal patterns in a wide range of biofilms, offering the potential for new insights into infection and pathogenicity.

## Introduction

Biofilms are biological structures formed by microorganisms in a variety of environments, and they consist of cells embedded in a self-secreted extracellular matrix [1]. Biofilm-forming organisms have evolved numerous mechanisms to protect their constituent cells from external biological and physiochemical stresses, resulting in populations of cells that exhibit increased resistance to a wide range of deleterious agents when compared to planktonic cells [2].

The structure of biofilms is integral to their role in infection, with mechanical deformation of soft substrates and epithelial cell monolayers being a major contributor to pathogenicity [3]. Pattern formation within biofilms is known to be affected by a change in the bulk carbon or nitrogen concentration of growth media [4, 5], as well as by nonuniform growth due to frictional forces and nutrient depletion [6]. Environmental and surface properties also affect growth parameters, leading to biofilms being a major burden to public health and industry [7]. It has been shown that varying substrate stiffness can alter cell attachment [8, 9], motility [10], growth dynamics [11] and expansion of constituent cells within biofilms [12], all of which are linked to infection.

It has long been known that bacterial growth rate and biomass formation depend on the type and concentration of nutrients available, and that altering the nature and availability of growth substrates can influence the growth of planktonic bacteria [13, 14, 15]. This has also been demonstrated for sessile biofilms, where growth substrates can have profound effects on biofilm morphology, growth dynamics, and mechanical properties [16, 17, 18, 19]. Recent studies on single-species biofilms have identified crossfeeding mechanisms for acetate, alanine and other nutrients influencing biofilm viability and morphology [20, 21, 22]. Emergent biofilm properties have also been found to give rise to water and nutrient transport mechanisms [23, 24, 25], yet the factors governing how they form and how environmental cues govern their structure are currently unknown.

Complex channel networks were recently discovered in *E. coli* by Rooney et al. [26] using the Mesolens, an optical mesoscope enabling sub-micron resolution across multi-millimeter size live biofilms [27]. These channels were found to transport small fluorescent microspheres, and their interior showed a higher accumulation of nutrients with respect to the rest of the biofilm. However, the large size of Mesolens datasets meant that only approximate measurements of intra-colony channel width could be made.

We hypothised that nutrient availability and agar concentration (a proxy for substrate stiffness) play an important role in channel morphology in mature *E. coli* biofilms, and we have developed a new open-source image analysis pipeline to test this conjecture. Our work shows that channel structure and distribution are strongly affected by both nutrient availability and substrate stiffness, and that the relative position of channels inside the biofilm determines their width for a given nutritional profile. The combination of Mesolens imaging and our image analysis pipeline provides precise and absolute measurements of intra-colony channel widths in response to specific environmental conditions and gives new insights into the role of nutrient availability in biofilm growth and sustenance.

## Results

### Nutrient availability affects biofilm morphology at the mesoscale

To test how biofilm morphology is affected by nutrient availability, specific carbon and nitrogen limiting conditions were determined for *E. coli* JM105-miniTn7-*gfp* (Supplementary Figure 1). Carbon and nitrogen were limited by controlling the nutrient concentrations of the growth media (glucose and ammonium chloride, respectively). The limiting carbon concentration was 15 ± 5 mM (C:N ratio of 1:1.25) and the limiting nitrogen concentration was 2.5 ± 1.5 mM (C:N ratio 27:1). These are broadly comparable with previous *E. coli* growth experiments in minimal medium [28].

We investigated the impact of nutrient concentration on biofilm morphology by measuring the global biofilm property of base area as a proxy for colony size and spreading (Supplementary Figure 2). Biofilms grown on glucose-limited substrates and those grown on ammonium-limited substrates were similar in size, whereas glucose excess led to larger biofilms than ammonium excess (p = 8.152 × 10^-5^). As expected, biofilms grown on glucose-rich media were larger than those grown under glucose limitation (p = 1.968 × 10^-5^). These data show that glucose availability is the limiting factor governing biofilm base area during growth on minimal medium.

Biofilm maximum intensity projection images (Figure 1) which are colour-coded by depth also indicate that not all biofilms have the uniform dome-shaped structure typical of *E. coli* biofilms [29, 30]. Instead, in Figure 1c we observe that the thickest region of the biofilm is located at an intermediate radius, between the centre and the edge of the biofilm. A phenomenon similar to colony sectoring is observed on glucose-limited substrates (Figure 1a), where radial sections of the biofilm have a considerably lower fluorescence signal than the rest of the biofilm. The pattern of intra-colony channels is also distributed heterogeneously inside the biofilm depending on the nutrient availability. Channels grown on nutrient-limited substrates appear to expand radially outwards from the centre in approximately straight lines, whereas on nutrient-rich substrates the channels often change direction sharply and do not follow a straight line. These observations confirm that the morphology of *E. coli* biofilms is strongly determined by nutrient availability.

**Figure 1:**
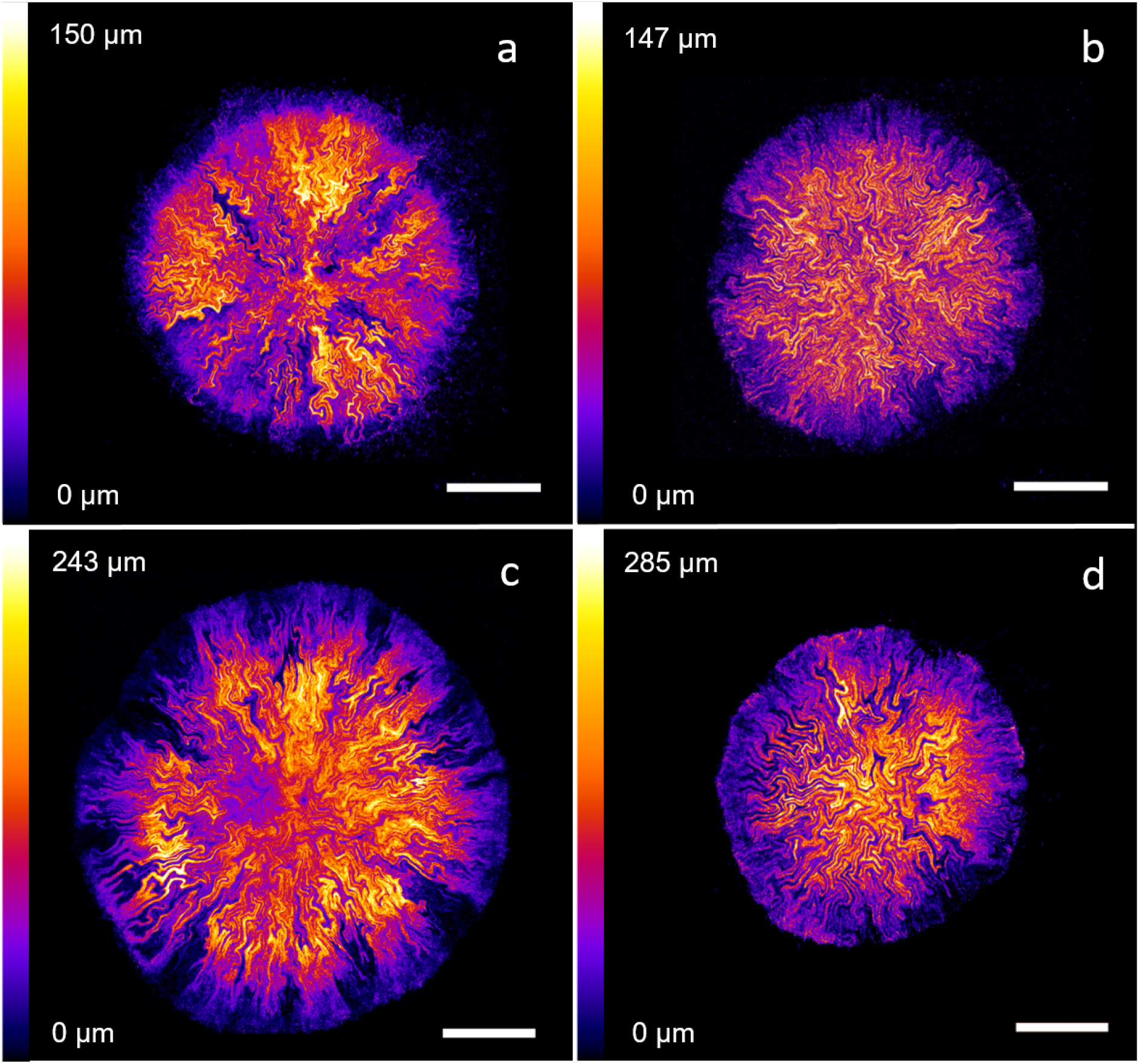
*E. coli* JM105-mini-Tn7-*gfp* biofilms grown on M9 minimal medium substrates with 15 mM carbon (a), 200 mM carbon (b), 2.5 mM nitrogen (c) and 10 mM nitrogen (d) concentrations. Intra-colony channel patterns appear radially expanding from the centre on nutrient-limited substrates (a,c), whereas they have a fractal structure with sharp turns on nutrient-sufficient substrates (b,d). The figure includes maximum intensity projections of confocal z-stacks made of images separated by 3 *μ*m in the z-direction. Images are colour-coded in the axial direction, with purple corresponding to the base of the biofilm and white corresponding to the top. Scale bars: 500 *μ*m.

### Intra-colony channel width increases non-linearly with radial distance from the centre of the biofilm

Channel width was measured along whole circumferences in *E. coli* JM105-miniTn7-*gfp* biofilms grown on M9 minimal medium substrates with limitation and excess of both glucose and ammonium chloride. The average channel width was plotted against radial distance from the centre of the biofilm, revealing a non-linear increase in width with radius. Representative plots of n = 3 repeats for each nutrient condition are shown in Figure 2. The minimum observed channel width was approximately 10 *μ*m, and corresponded to the innermost region of the biofilm (at 200 *μ*m radius). This value was not limited by the resolution of the image datasets, but rather by the smallest radial distance at which channel width could be accurately measured by our pipeline.

**Figure 2:**
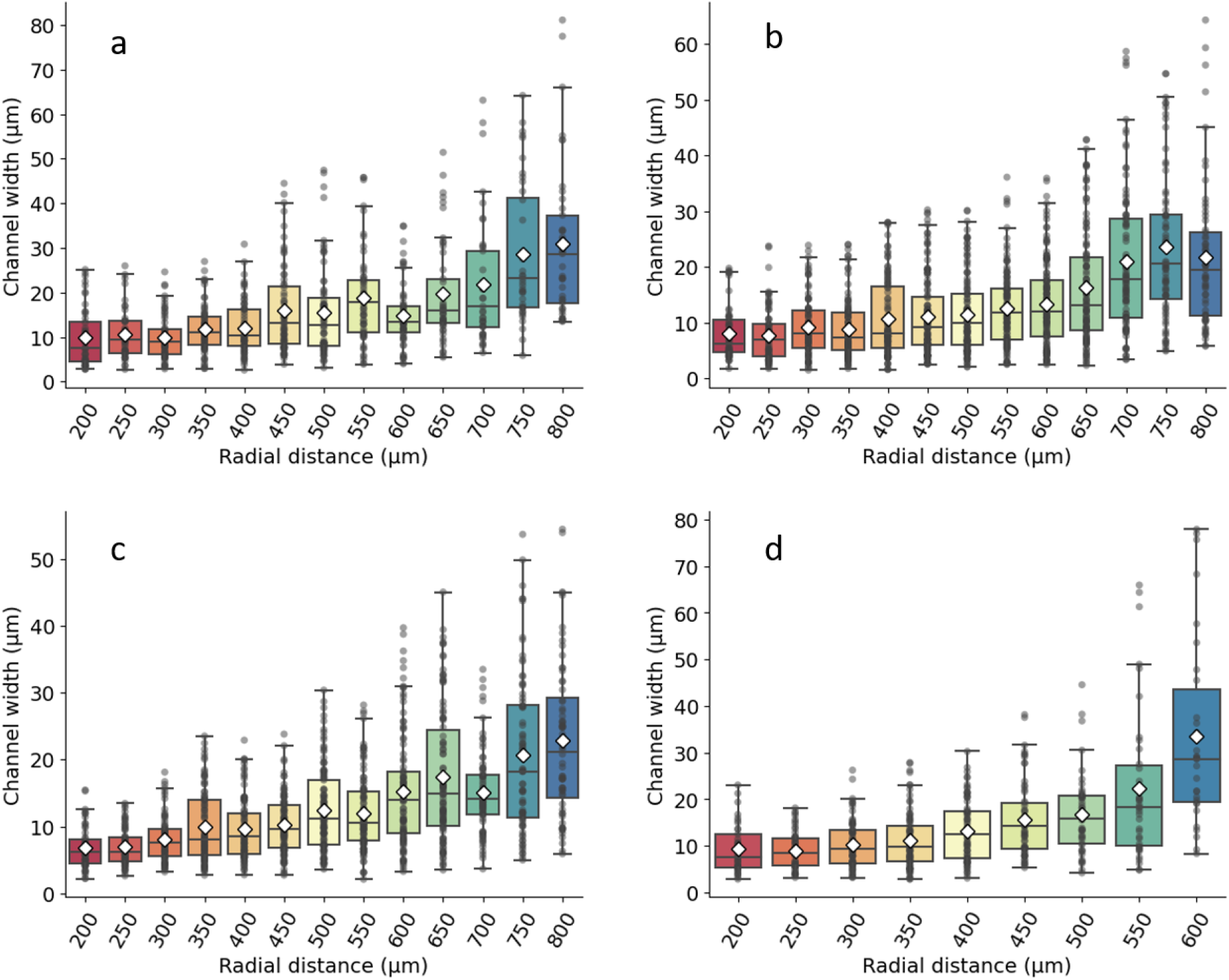
Non-linear channel width variation across the radial direction for *E. coli* JM105-miniTn7-*gfp* biofilms grown on M9 minimal medium substrates with 15 mM carbon (a), 200 mM carbon (b), 2.5 mM nitrogen (c) and 10 mM nitrogen (d). Each plot is representative of n = 3 biological repeats. Average values are shown as white diamonds, whereas boxes represent the interquartile range (with median values shown as horizontal lines inside each box).

The modality of increase in intra-colony channel width was explored by applying linear and exponential fits to each dataset, and R-squared values were calculated for each fit. Average R-squared values for exponential fits were 0.845 for 15 mM carbon, 0.880 for 200 mM carbon, 0.882 for 2.5 mM nitrogen and 0.878920 for 10 mM nitrogen. By verifying that the increase in channel width along the biofilm radius was non-linear, we ensured that the change was not just a result of polar geometry.

### Glucose-limited biofilms possess wider channels than ammonium-limited biofilms

To investigate the effect of nutrient availability on intra-colony channel width, colonial biofilms were grown on substrates with a range of nutrient concentrations: 15 mM carbon (n = 8 biofilms), 200 mM carbon (n = 7 biofilms), 2.5 mM nitrogen (n = 9 biofilms) and 10 mM nitrogen (n = 4 biofilms). Intra-colony channel width was compared across nutrient conditions at three normalised radial positions (Supplementary Figure 4), revealing that channels were approximately 25% wider on glucose-limited substrates than on ammonium-limited substrates inside the biofilm. This increase was most significant at the mid-radius region of each biofilm, where channel widths measured on average 13.78 *μ*m under glucose limitation and 11.27 *μ*m under ammonium limitation.

These data suggest that carbon- and nitrogen-based nutrients mechanistically affect channel morphology in different manners. The density of channels detected inside the biofilm also varied differently depending on the nutrient source, and was largest at mid-radius.

### Substrate stiffness affects channel density and biofilm base area

The effect of substrate stiffness (determined by agar concentration) on internal biofilm morphology was investigated by imaging biofilms grown on soft and hard agar substrates, in both rich and minimal medium (Figure 3). We observed an increase in out-of-focus fluorescence for biofilms grown on rich medium substrates compared to biofilms grown on minimal medium substrates. On minimal medium, channel borders were better resolved due to the higher contrast relative to the rest of the biofilm. On rich medium, the density of intra-colony channels increased with decreasing substrate stiffness. On 0.5% agar LB substrates (Figure 3a) channels were densely packed in the whole biofilm, while some widely-separated channels progressively appeared on portions of biofilms grown on stiffer substrates (1% and 2% agar concentration, Figures 3b and 3c respectively).

**Figure 3:**
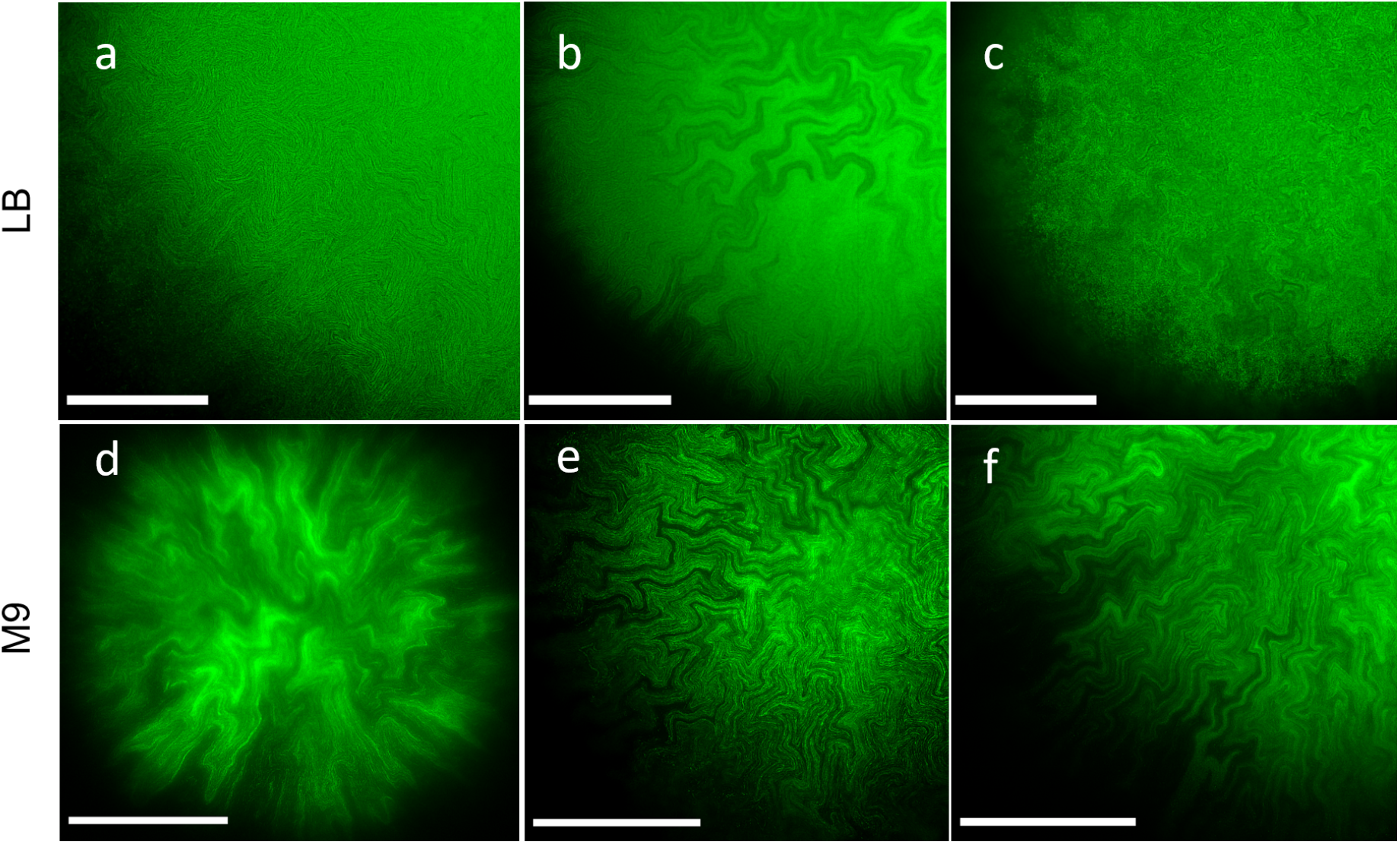
ROIs showing intra-colony channel morphologies of *E. coli* JM105-mini-Tn7-*gfp* biofilms grown on solid LB rich medium (top) and solid M9 minimal medium (bottom). The agar concentrations of the substrates were 0.5% (a,d), 1% (b,e) and 2% (c,f). All images were acquired using the Mesolens in widefield epi-fluorescence mode to capture the biofilms at the midpoint along biofilm thickness. The images were acquired with LB or M9 mounting medium and the Mesolens in water immersion, except for panel d, which was acquired in air with no mounting liquid to prevent the biofilm from detaching from the substrate. All images were deconvolved using Huygens proprietary software. Scale bars: 500 *μ*m.

Varying agar concentration was also associated with a change in biofilm base area for both rich and minimal medium substrates (Supplementary Figure 3). On rich LB medium substrates, decreasing agar concentration resulted in an increase in biofilm base area, whereas the effect was opposite on M9 minimal medium substrates. Mann-Whitney U rank tests were performed between the 0.5% and the 2% agar concentration data for both rich and minimal medium, and revealed that increasing agar stiffness had similar effects on biofilm base area for minimal medium (p = 0.0221) and rich medium (p = 0.0180). However, increasing agar concentration led to an increase in biofilm base area for minimal medium, and a reduction for rich medium. The difference in base area between biofilms grown on rich and minimal medium substrates was most significant at 0.5% agar concentration (p = 0.0092).

## Discussion

This study constitutes the first quantitative analysis of intra-colony channels in mature *E. coli* biofilms. Using a simple custom image analysis pipeline based on the open-source software FIJI and the Python programming language, it was possible to quantify intra-colony channel width subject to various environmental conditions. We discovered that channel morphology was affected by both the type of substrate (rich or minimal medium agar) and by substrate properties (nutrient and agar concentration).

Environmental conditions altered the distribution of intra-colony channels inside biofilms. Channels formed on substrates with limiting carbon or nitrogen concentrations expanded radially outwards, in approximately straight lines. This could be because bacteria growing on nutrient-depleted substrates adhere less strongly on the surface of attachment, and expand more rapidly [31], or may be indicative of nutrient foraging behaviour. Conversely, channels formed on nutrient-rich substrates exhibited complex fractal patterns. It is hypothesised that this type of structure emerges due to rapid cell growth and larger cell dimensions, thereby resulting in a tighter network of channels. This in turn enables a greater proportion of cells to access the nutrients that are transported within the biofilm. A recent study by Fei et al. [6] highlighted two distinct types of patterns forming on the surface of *V. cholerae* biofilms, namely radial stripes and zigzag herringbone patterns. Conversely, the biofilms in our study only involved the same type of fractal patterns: intra-colony channel structures had homogeneous morphology, with only width and spacing varying depending on environmental conditions.

Furthermore, we found that agar concentration in the substrate affected intra-colony channel density on rich medium substrates: channels were tightly packed and had a uniform, narrow width on semisolid substrates, but as the agar concentration increased, wider channels appeared in portions of the biofilm. This observation appears to be in contrast with previous work on *V. cholerae* biofilms, where homogeneity in radial feature distribution and spacing at the edge of mature biofilms increased with agar stiffness [32].

An increase in out-of-focus fluorescence was also observed in widefield epi-fluorescence images of biofilms grown on rich medium compared to those grown on minimal medium. The fluorescence signal inside channels was likely due to bacterial cells being present inside them, as a result of movement inside the biofilm, collective reorientation [33] or shedding of cells from the inner walls of the channels. Previous studies on *V. cholerae* [34] confirmed that expansion in the depth direction is dependent on friction between the substrate and the expanding cells, with softer substrates leading to flatter biofilms than stiff substrates. This could explain the reduction in contrast between channel border and constituent cells observed on stiff rich medium substrates. A variation in internal biofilm architecture along the depth direction has also previously been reported for *E. coli* growing on agar substrates [35] and was attributed to differences in extracellular matrix components assembly and organisation inside the biofilm volume.

The variation in channel width inside mature biofilms was discovered to increase non-linearly along the radial direction from the centre of the biofilm under all nutrient conditions. Remarkably, channels were on average 25% wider at the centre of glucose-limited media than inside ammonium-limited media, which suggests that they are more important for the transport of carbon-based nutrients. We speculate that the increase in channel width at the edge of the biofilm is due to rapid cell growth in nutrient-rich conditions [36]. This would result in non-uniform radial expansion, as previously observed for *B. subtilis* [31] and *V. cholerae* [6], suggesting that variations in channel width are an emergent property of *E. coli* biofilm growth.

We speculate that the channels may become increasingly narrow in the centre of the biofilm because of fundamental fluid dynamics behaviour. We can consider a simplified model of the nutrients transported by the channels in an *E. coli* biofilm as fluid flow in a pipe. The continuity equation states that in the case of steady flow the mass flow rate remains constant: as fluid flows along a pipe of decreasing width, the velocity of the fluid increases proportionally [37]. Approximating our channels as pipes, the smaller width of the channels at the centre of the biofilm compared to the edge will result in a faster provision of nutrients where the demand is greatest to sustain the biofilm. In addition, narrow channels may also help to transport nutrients into the biofilm centre via capillary action. The distance over which fluid can be spontaneously transported through capillary action is inversely proportional to the width of a pipe [38]: the smaller channel width towards the centre of the biofilm would then support increased capillary action and hence provide an additional nutrient transport mechanism.

The mesoscopic effects of nutrient limitation were studied by calculating biofilm substrate area. Nutrient abundance in the substrate, achieved by increasing the amounts of either glucose or ammonia, led to larger biofilms than nutrient limitation. Biofilm base area was most significantly affected by the glucose concentration in the growth substrate and was 2.7 times larger for biofilms grown on glucose excess than on glucose limitation. On the other hand, increasing the ammonium concentration on a substrate with sufficient glucose availability did not significantly affect biofilm base area. These findings agree with the reported increase of biofilm formation with glucose concentration in *Staphylococcus* species [39] and the increase in colony size with glucose concentration for *E. coli* [40] and yeast [41]. While it is known that growth rate limitations and carrying capacity limitations differ due to the change in growth condition (homogeneous mixed environment or growth on a solid surface) [42], our data show a similar response to both conditions (Supplementary Figures 1 and 2).

The effect of agar concentration on biofilm base area varied based on the nutritional profile of the substrate. An increase in agar concentration in rich medium was associated with a reduction in biofilm base area, due to slower radial expansion of biofilms on the surface brought by stronger frictional forces [29]. An increase in agar concentration could also affect biofilm base area by decreasing the level of swarming motility [43]. The trend of increased base area with decreased agar stiffness has previously been observed in *Vibrio cholerae* biofilms grown on LB medium with different agar concentrations [6]. However, in our work the opposite trend was observed on minimal medium: biofilms grown on stiff substrates had a base area on average 1.6 times larger than those grown on semi-solid substrates. This could be due to faster assimilation of nutrients occurring on semi-solid minimal media with respect to stiff minimal media, causing a halt to radial expansion. Finally, biofilm base area was on average almost 2.5 times larger on semi-solid rich medium substrates than on semi-solid minimal medium substrates. The slower growth of *E. coli* in minimal medium with respect to rich medium (with average doubling times of 18 ± 0.5 minutes in LB and 38 ± 1.1 minutes in M9-glucose [44]) can explain this difference in biofilm base area only in part, since we observed no significant difference in base area between rich and minimal medium substrates at higher agar concentrations of 1% and 2%.

Studying biofilm internal patterns at the cellular level is a task mostly relegated to theoretical modelling and computer simulations [29, 45, 46, 47, 48]. Experimental approaches in imaging biofilms at single-cell resolution [49, 50, 51] have been limited in the total biofilm size that can be imaged at once (under 100 *μ*m^2^). The combination of mesoscale imaging and open-source image analysis tools has proven to be an essential platform technology to identify and quantify internal structure in larger biofilms than an ordinary microscope can accommodate. It can also be used without adaptation to validate existing numerical simulations of mature colony biofilm structure, or it can be readily applied for the quantification of any internal patterns and structures in a wide range of biofilms.

## Methods

### Materials and strains

The non-pathogenic *E. coli* strain JM105 (endA1, recA1, gyrA96, thi, hsdR17 (rk–, mk+), relA1, supE44, Δ(lac-proAB), [F’ traD36, proAB, laqIqZΔM15]) containing mini-T7 *gfp* [52], enabling *gfp* fluorescence, was used throughout the study. Liquid cultures were grown in Lysogeny Broth (LB) [53]. Minimal medium growth was in M9 medium [53], prepared in a 5x concentrated solution (Na_2_HPO_4_ 30 g/L, KH2PO4 15 g/L, NH_4_Cl 5 g/L, NaCl 2.5 g/L) then diluted to 1x with distilled deionised water and supplemented with 1 mM MgSO_4_· 7H_2_O, 0.2% (w/v) glucose and 0.00005% (w/v) thiamine. Solid substrates were made by adding agar prior to autoclaving, in concentrations of 20 g/L (LB agar) and 15 g/L (M9 agar) for nutrient concentration experiments. For agar stiffness experiments, LB and M9 media were prepared as described above, but the agar concentration was progressively reduced for soft substrates (giving 5 g/L, 10 g/L and 20 g/L for both LB and M9 media).

### Imaging chamber design and 3D-printing

The imaging chamber to be used with the Mesolens was designed on AutoCAD (Autodesk, USA) by modifying the original design of Rooney et al. [26]. The chamber was designed to mimic a Petri dish, and consisted of a 120 mm x 100 mm x 12 mm plate with a 60 mm diameter, 10 mm deep well at its centre to hold the agar substrate (Supplementary Figure 5). After 3D printing the chamber in black ABS plastic (FlashForge, Hong Kong) using a FlashForge Dreamer 3D printer (FlashForge, Hong Kong), the corners were smoothed out using a scalpel. This reduced the movement of the imaging chamber inside the square plate. The chamber was sterilised with a 70% (v/v) ethanol solution, and then under UV light for 20 minutes immediately before the addition of sterile solid growth medium.

### *E. coli* biofilm specimen preparation

Liquid cultures of *E. coli* JM105-mini-Tn7-*gfp* were prepared in LB medium supplemented with 25 *μ*g/mL gentamicin and incubated overnight at 37 °C. Overnight cultures were diluted (1:100) into fresh LB medium and incubated at 37 °C until they reached an optical density of 0.5 (mid-exponential growth phase).

To grow biofilms on rich medium, 100 *μ*L of a liquid culture with density 1×10^4^ colony-forming units (CFUs) per mL was inoculated on the 10 mm-thick LB agar substrate inside the imaging chamber. The specimen was incubated at 37 ^°^C for 24 hours in darkened condition prior to imaging. For minimal medium with varying nutrient concentration, mid-exponential phase liquid cultures were washed three times with 1x M9 salts. They were then resuspended in M9 medium with appropriate amounts of glucose and ammonium chloride (for carbon and nitrogen variation, respectively). M9 agar substrates cast into the 3D-printed imaging chambers were inoculated at a concentration of 1×10^4^/mL. Based on the dimensions of the imaging chamber (Supplementary Figure 5), this corresponds to a seeding density of 1 cell per mm^2^, which ensured biofilms were sufficiently spaced out and did not have to compete for nutrients with others in their proximity. It was also ensured that only one biofilm was visible in the field of view of the Mesolens at once, which prevented background signal from nearby biofilms from reaching the detectors.

### *E. coli* growth characterisation under different nutrient conditions

Mid-exponential growth phase liquid cultures of *E. coli* JM105-mini-Tn7-*gfp* were prepared as described above, then washed and resuspended in 1x M9 salts. The cultures were diluted to an OD_600_ of 0.04, split in individual tubes and supplemented with appropriate amounts of the nutrient of interest. For carbon growth curves, the concentration of nitrogen was kept constant at 18.7 mM, and for nitrogen growth curves the concentration of carbon was kept constant at 66.7 mM. The nutrient concentrations were chosen to be between 0 mM and 80 mM - for comparison, the carbon and nitrogen concentrations in nominal M9 salts are 66.6 mM and 18.7 mM, respectively. Aliquots (200 *μ*L) of liquid culture of each investigated concentration was added in triplicate to a black Nunc MicroWell 96-well optical-bottom plate with polymer base (ThermoFisher Scientific, US). Absorbance (OD_600_) measurements were performed every 15 minutes for 24 hours, with the plate being held at 37° C and shaken continuously.

Growth curves were produced for each nutrient concentration thanks to the Gen5 microplate software (BioTek, USA), and exported to MATLAB (MathWorks, USA) for analysis. The average of the first absorbance value for each concentration was used as a baseline and subtracted from each growth curve. The y axis was displayed in logarithmic scale, in order to identify the exponential growth phase (the linear portion of the plot). A linear fit to this region was applied using MATLAB’s Curve Fitting Toolbox by selecting an exponential function of the form *y* = *a exp(bx),* with the specific growth rate corresponding to the coefficient b. While error analysis was produced internally in MATLAB as R-squared, SSE and RMSE values, the error bars on the specific growth rate plots were calculated as the standard deviations across biological repeats for the same nutrient concentration. Note that the duration of the exponential growth condition varied depending on the nutrient concentration, and lasted between 6 and 18 hours, hence the data points corresponding to exponential growth were selected manually. Growth curves and specific growth rates were plotted in Python using matplotlib.

### Widefield epi-fluorescence and confocal laser scanning Mesolens imaging

All the data in this work was acquired using the Mesolens, a custom-made optical microscope combining high numerical aperture (0.47) with low magnification (4x). These characteristics allow simultaneous imaging over a field of view of 6 mm x 6 mm with sub-cellular lateral resolution throughout (700 nm). Images can be acquired over a total thickness of 3 mm with an axial resolution of 7 *μ*m. For widefield epi-fluorescence Mesolens imaging, GFP fluorescence was excited by a 490 nm pE-4000 LED (CoolLED, UK), and emitted fluorescent light was made to pass through a 540 ± 10 nm bandpass filter before being detected by a VNP-29MC sensor-shifting CCD camera (Vieworks, South Korea). The camera uses sensor-shifting to acquire 9 images for each pixel (each shifted by 1/3 of a pixel width from the other, forming a square grid), effectively increasing the resolution to 259.5 MP [54]. In confocal laser scanning mode, GFP fluorescence was excited using a 488 nm laser (Multiline Laserbank, Cairn) at 5mW power. Fluorescence emission was filtered through a 525/39 nm bandpass filter (MF525-39, Thorlabs, USA) before being detected using a PMT (PMM02, Thorlabs, USA). The emission path also included a 505 nm longpass dichroic mirror (DMLP505R, Thorlabs, USA). Both widefield and confocal laser scanning microscopy were performed with the lens in water immersion, to match the refractive index of the mounting media (LB and M9), except the 0.5% M9 agar datasets which were acquired in air, with no immersion or mounting liquid. This was needed to preserve biofilm structure, as biofilms grown 0.5% M9 agar were less adherent to the substrate and detached from it after addition of liquid mounting medium. The intra-colony channels formed by these biofilms remain evident even though the lower refractive index resulted in poorer spatial resolution (Figure 3d). The Mesolens collars were adjusted to minimise spherical aberrations.

### Image analysis

Confocal z-stacks were displayed as a maximum intensity projection, colour coded by depth across the z-direction using the “Fire” lookup table from FIJI (ImageJ version 1.53c). Where necessary, median-filtered widefield epi-fluorescence Mesolens images (filter radius: 2 pixels) were deconvoluted using the proprietary Huygens Professional version 19.04 software (Scientific Volume Imaging, Netherlands). Deconvolution was performed after an in-built manual background subtraction and a theoretical point spread function estimation on Huygens, using the Classic Maximum Likelihood Estimation method with 50 iterations, a signal-to-noise ratio of 40 and a quality threshold of 0.01.

To calculate biofilm base area, widefield Mesolens images were opened in FIJI, and thresholded using the “Adjust Threshold” tool, using the mean of gray levels as the threshold value. The “Wand (tracing) tool” was then used to select the biofilm mask, and the area was calculated using the “Measure” function. The base area measurements were systematically underestimated due to the thresholding method used by FIJI to create the binary mask. Nonetheless, by using the same thresholding method on all images we ensured that this limitation had comparable effects on all datasets. Biofilm thickness was calculated by plotting z-axis profiles of confocal z-stacks in FIJI and calculating the distance between the two minima on each profile.

Mesolens images were brightness- and contrast-adjusted using FIJI for presentation purposes [55].

### Intra-colony channel width calculation

Intra-colony channel width was calculated using the image analysis workflow outlined in Figure 4. The pipeline made use of the FIJI plugin “Polar transformer” [56], which performed an image transformation from polar to cartesian coordinates, as well as of two custom Python scripts. Initially, the coordinates of the centre of the biofilm were calculated using the “Measure” function on FIJI applied on an oval selection of the whole biofilm. The Polar Transformer plugin was then launched, and the origin of transformation was manually entered as the coordinates of the biofilm’s centre. The number of pixels for each degree in angular coordinates was set to 7200 (corresponding to 20 pixels for each degree). The polar-transformed image was contrast-adjusted using the “CLAHE” (Contrast Limited Adaptive Histogram Equalization) feature on FIJI with block size 60, maximum slope 3 and 256 histogram bins to facilitate channel identification. The image was then despeckled using the “Despeckle” function on FIJI in order to remove noise. The image was also inverted, making intra-colony channels appear light and cells dark, which facilitated the rest of the analysis. Vertical line selections were taken at different x positions on the polar-transformed image, corresponding to circumferences around the biofilm taken at different radial distances from its centre. Signal profiles were obtained for each line selection using the “Plot profile” feature on FIJI, and were exported to Python, where signal peaks were located thanks to a custom script using the find\_peaks() function. For each dataset, peak thresholding was performed in order to exclude noise: peak prominence was chosen as 20% of the difference between maximum and minimum signal, and a minimum distance between adjacent peaks was selected as 9 pixels, which corresponds to the average length of an *E. coli* cell (2 *μ*m) on a Mesolens widefield epifluorescence image. Intra-colony channel widths were then calculated as the full-width at half-maximum of each peak. The actual width of the channels in *μ*m was converted according to the radial distance from the centre, using polar coordinate geometry.

**Figure 4:**
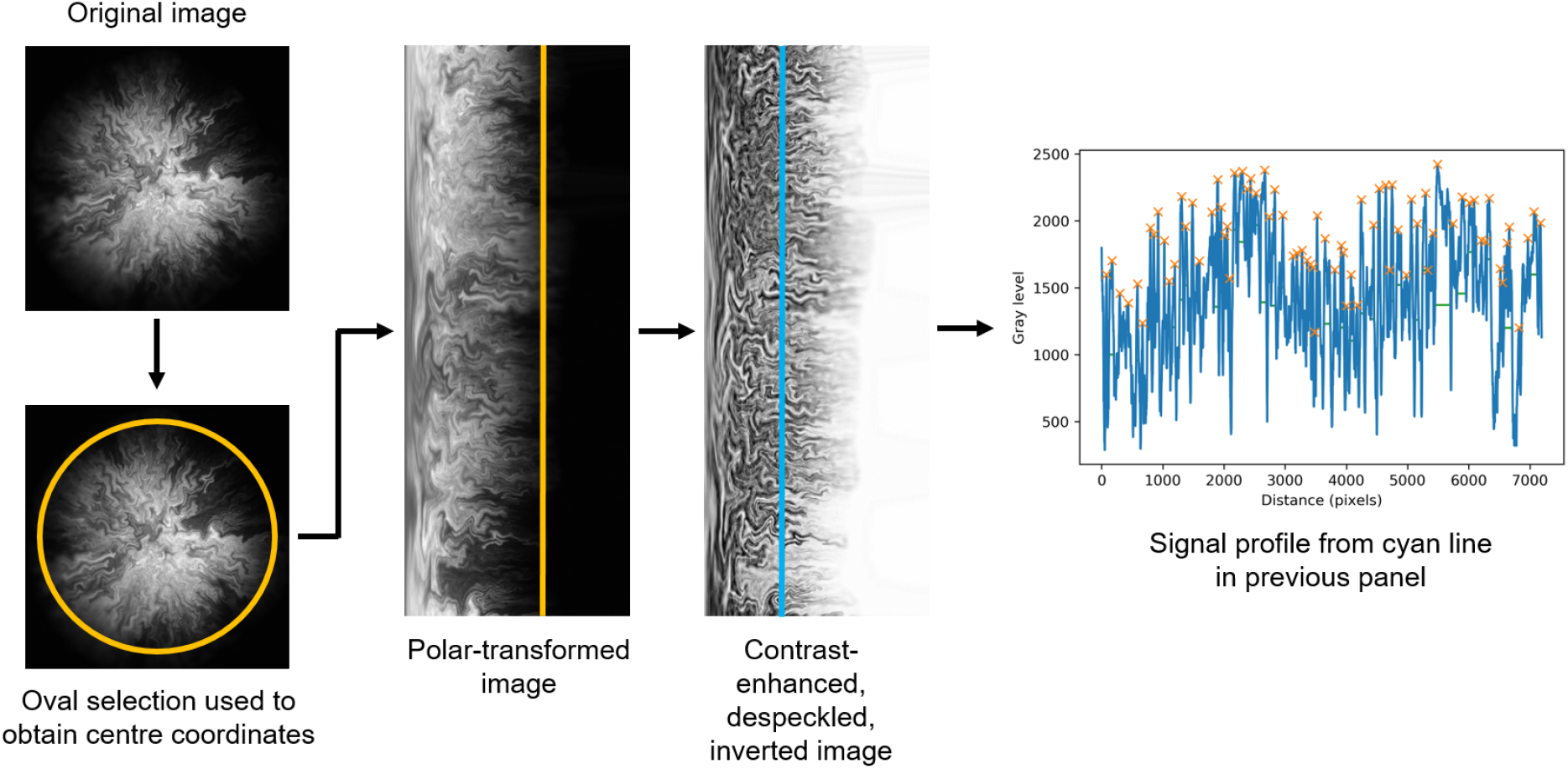
Image analysis workflow. A widefield Mesolens image is opened on FIJI, where an oval selection is used to obtain the coordinates of the centre of the biofilm. The coordinates are the input into the Polar Transformer FIJI plugin, which performs a transformation from polar to cartesian coordinates. The resulting image is locally contrast-enhanced, despeckled and inverted on FIJI. Signal profiles of vertical line selections are exported to Python, where the signal is analysed with a custom script in order to locate the peaks (orange crosses) and calculate the full-width at half-maximum (green horizontal lines) of each peak. The latter quantity corresponds to intra-colony channel width, converted from pixel units to *μ*m using polar geometry.

### Variation of intra-colony channel width along the radial dimension

Intra-colony channel width was measured along whole circumferences at radius intervals of 50 *μ*m, starting from an initial radius of 200 *μ*m. This initial radius value was chosen because at lower values, polar transformed images were distorted and signal analysis was not reliable. Because of the ellipticity of the biofilms, polar-transformed images appeared as rectangles with a non-straight right side (Figure 4), corresponding to the biofilm perimeter. Regions beyond which a full circumference could not be obtained for the biofilm were excluded from the analysis.

Channel widths were measured for biofilms grown on the four nutrient conditions described in the Results section. Three biological replicates were used for each condition, giving a total sample size of 12 biofilms. Linear and exponential fits were applied to each plot of average channel width against radius, and R-squared values for the fits were computed for each dataset using Excel. The number of channels identified by the script varied between radial positions, and was highest for the intermediate radii. This is likely due to the fractal nature of intra-colony channels, which have sharp changes in direction and fold on themselves inside the biofilm.

To compare channel widths across different nutrient concentrations of the substrate, intra-colony channel width was calculated at three normalised radial positions (full radius, 50% radius and 20% radius) to investigate the effect of nutrient concentration. Datasets were acquired for each of the following nutrient concentrations: 15 mM carbon (8 biofilms, giving 1062 channels), 200 mM carbon (7 biofilms, giving 1424 channels), 2.5 mM nitrogen (9 biofilms, giving 1524 channels) and 10 mM nitrogen (4 biofilms, giving 507 channels).

### Statistical analysis

The comparison of biofilm base areas and channel widths under different nutrient availability and agar concentration was performed by means of a Mann-Whitney U rank test, chosen as datasets were not normally distributed and had unequal sample sizes. The tests were performed using the `scipy.stats.mannwhitneyu()` function in Python. P-values smaller than 0.05 were considered statistically significant. Biofilm base area and intra-colony channel measurements were displayed using Python’s `seaborn.boxplot()` and `seaborn.stripplot()` plotting tools, where the first and third quartile of the data (Q1 and Q3) are enclosed by a box which contains 50% of the data (the interquartile range, or IQR). The median is shown as a horizontal line inside each box, whereas the capped bars are the “minimum” and maximum” values, which are found a distance of 1.5*IQR above and below the Q3 and Q1 respectively. Average values for each measurement are shown as white diamonds. An outlier removal algorithm was used on the data to discard channels whose large width was attributed to colony sectoring. Outliers were removed by excluding data points which had a modified z-score greater than 3. The modified z-score was calculated using the median absolute deviation as described in [57] for non-normally distributed data.

## Data and code availability

The Python script used to calculate intra-colony channel width was written on Python version 3.7, and is deposited on GitHub (https://github.com/beatricebottura/biofilm_channel_width, DOI: 10.5281/zenodo.5786305). An example dataset to be analysed with the script is also available in the same repository. The remaining data used to generate channel width results is available upon reasonable request.

## Acknowledgements

We thank Dr Ainsley Beaton (John Innes Centre, formerly University of Strathclyde) for the kind gift of the JM105-miniTn7-*gfp* strain. We also acknowledge Dr John Munnoch (University of Strathclyde), Dr Rebecca McHugh (University of Glasgow), Dr Morgan Feeney (University of Strathclyde) and Jordan Murray for helpful discussions. Beatrice Bottura is supported by a University of Strathclyde Student Excellence Award. Gail McConnell is supported by the Medical Research Council [MR/K015583/1] and Biotechnology & Biological Sciences Research Council [BB/P02565X/1, BBT011602]. Paul A. Hoskisson is supported by funding from the Biotechnology & Biological Sciences Research Council [BB/T001038/1 and BB/T004126/1] and the Royal Academy of Engineering Research Chair Scheme for long term personal research support [RCSRF2021\11\15].

## Author Contributions

**Beatrice Bottura:** Conceptualisation, Methodology, Software, Validation, Formal analysis, Investigation, Data Curation, Writing - Original Draft, Writing - Review & Editing, Visualisation. **Liam Rooney:** Conceptualisation, Methodology, Writing - Review & Editing. **Paul A. Hoskisson:** Conceptualisation, Methodology, Formal analysis, Writing - Review & Editing, Resources, Supervision, Project administration, Funding acquisition. **Gail McConnell:** Conceptualisation, Methodology, Formal analysis, Writing - Review & Editing, Resources, Supervision, Project administration, Funding acquisition.

## Additional information

**Supplementary Figure 1:**
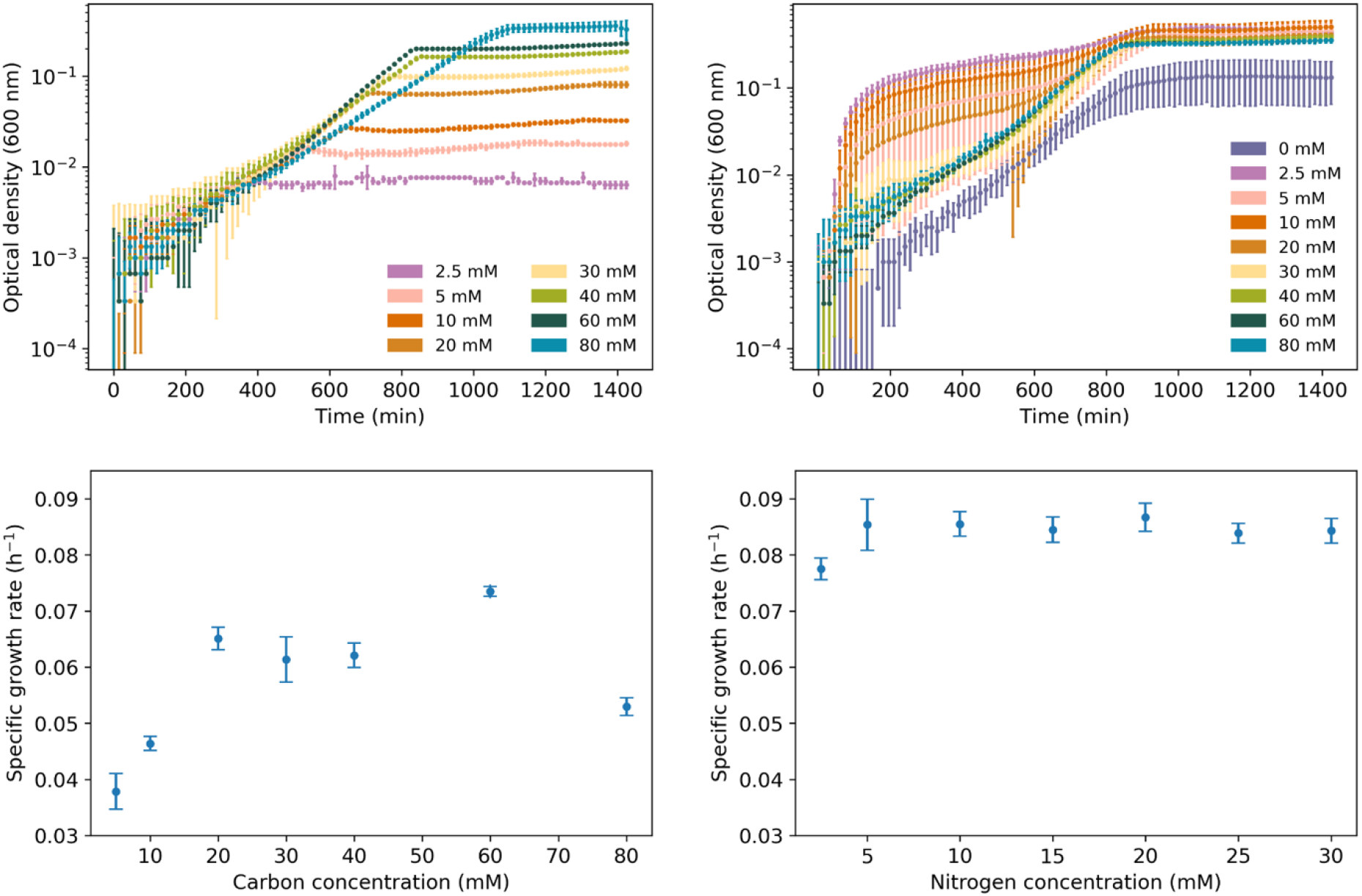
Growth curves (top panels) and specific growth rates (bottom panels) of *E. coli* JM105-mini-Tn7-*gfp* liquid cultures grown in M9 minimal medium with various carbon (a, c) and nitrogen (b, d) concentrations, obtained by varying the amounts of glucose and ammonium chloride in the medium. Error bars on growth curve plots represent the standard deviation across three biological repeats. The growth curve for the lowest carbon concentration in a (0 mM) is not shown as it consists of a baseline of non-growing cells. The data point at 80 mM carbon (c) is likely due to saturation, especially if we consider that the nominal carbon concentration in M9 medium is 67 mM.

**Supplementary Figure 2:**
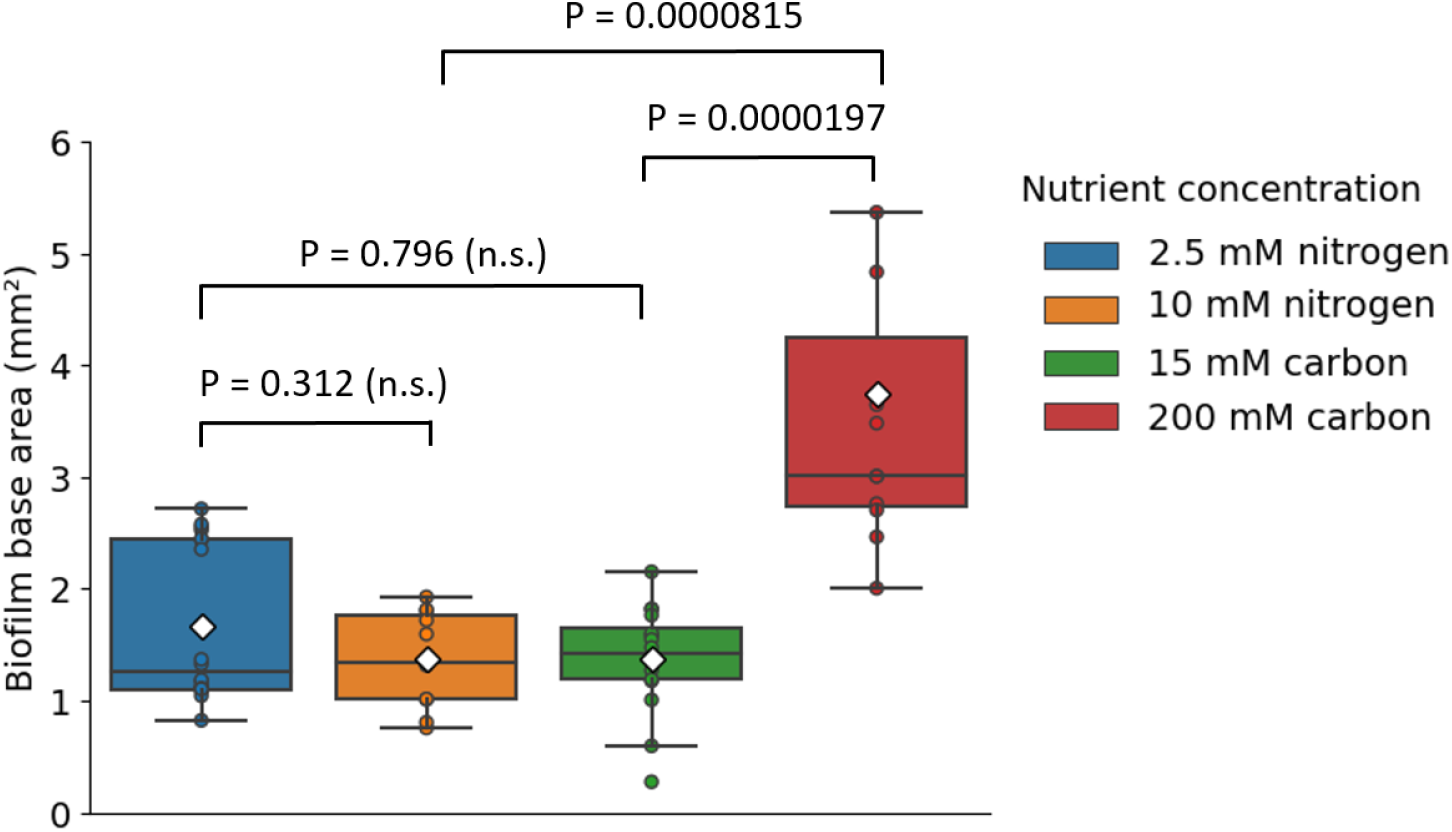
Base area of *E. coli* JM105-mini-Tn7-*gfp* biofilms calculated for four nutrient conditions. Average biofilm areas are 1.374 ± 0.118 mm^2^ (glucose-limited biofilms, n = 16), 3.738 ± 0.510 mm^2^ (glucose-rich biofilms, n = 11), 1.668 ± 0.166 mm^2^ (ammonium-limited biofilms, n = 18) and 1.370 ± 0.129 mm^2^ (ammonium-rich biofilms, n = 11). Uncertainties correspond to standard errors of the mean across biological repeats (n ≥ 11 for each condition). Mann-Whitney U rank tests were performed on the data, with relevant p-values shown. Average values are shown as white diamonds, whereas boxes represent the interquartile range (with median values shown as horizontal lines inside each box).

**Supplementary Figure 3:**
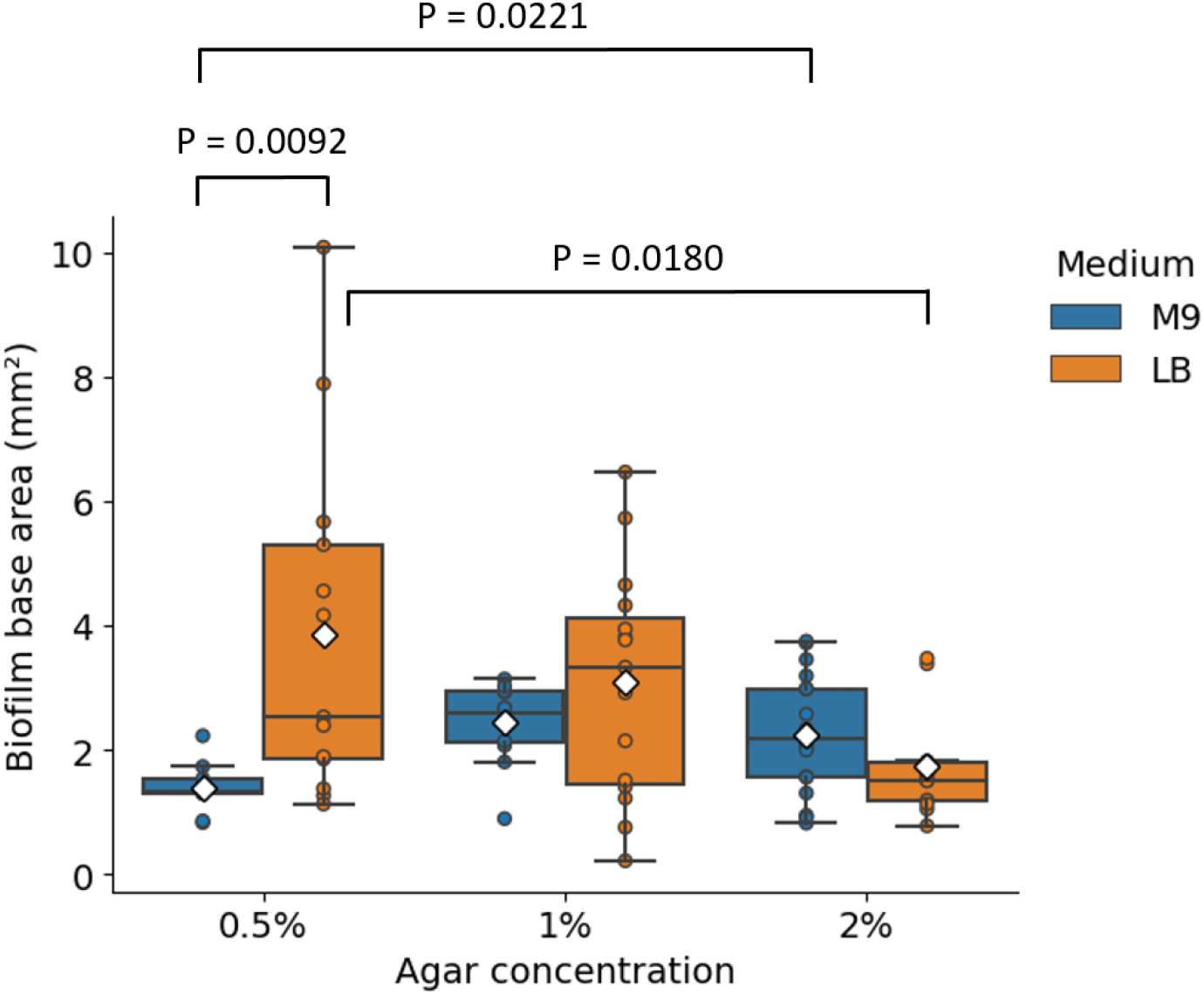
*E. coli* JM105-mini-Tn7-*gfp* biofilm base area calculated for three different agar concentrations of the substrate, in both rich (LB) and minimal (M9) medium. Average areas of biofilms grown on rich medium are 3.854 ± 0.772 mm^2^ (0.5% agar, n = 13 biofilms), 3.080 ± 0.477 mm^2^ (1% agar, n = 15 biofilms) and 1.733 ± 0.245 mm^2^ (2% agar, n = 12 biofilms). Average areas of biofilms grown on minimal medium are 1.387 ± 0.142 mm^2^ (0.5% agar, n = 9 biofilms), 2.434 ± 0.184 mm^2^ (1% agar, n = 12 biofilms) and 2.253 ± 0.224 (2% agar, n = 18 biofilms). Uncertainties correspond to standard errors on the mean across biological repeats. Mann-Whitney U rank tests were performed on the data, with relevant p-values shown. Average values are shown as white diamonds, whereas boxes represent the interquartile range (with median values shown as horizontal lines inside each box).

**Supplementary Figure 4:**
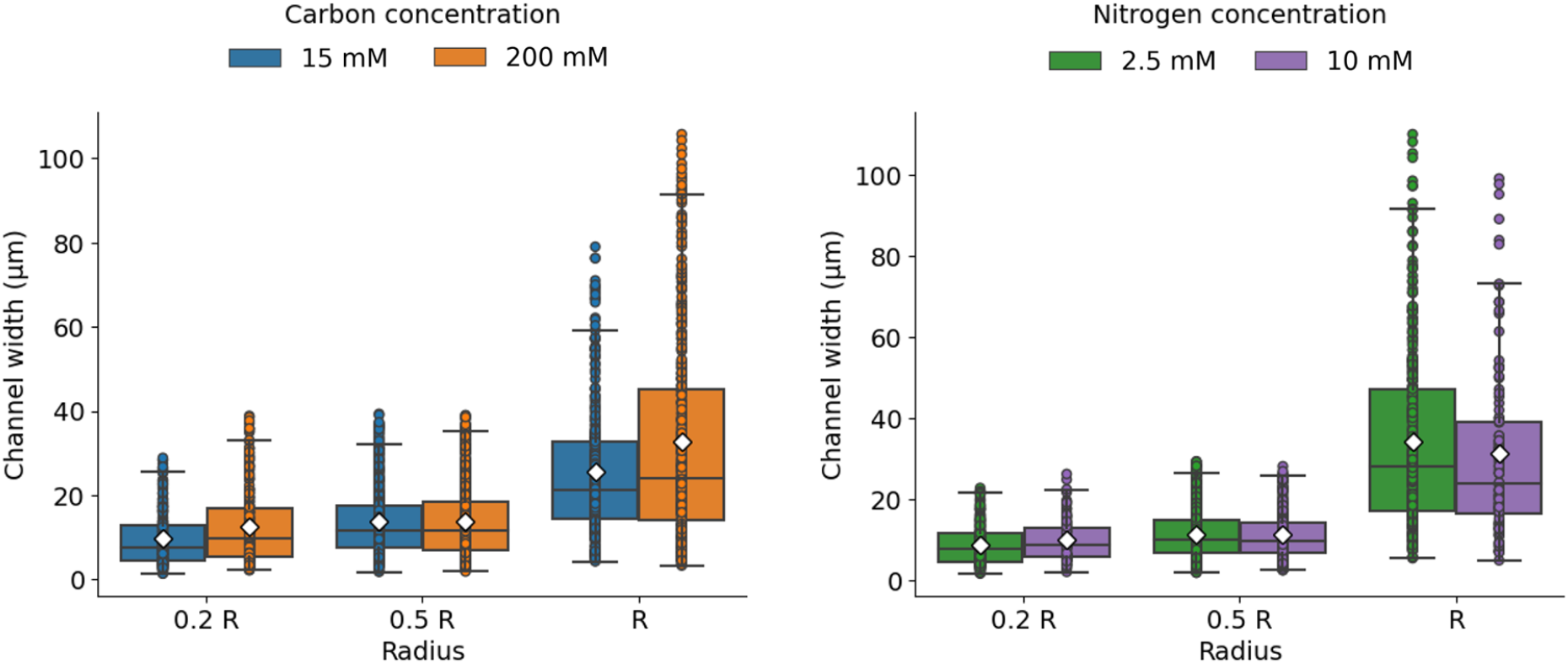
Intra-colony channel width calculated at three normalised radial positions (20% radius, 50% radius and full radius, labelled as 0.2 R, 0.5 R and R respectively). Channels were approximately 50% wider on glucose-limited substrates than on ammonium-limited substrates at the mid-radius region of each biofilm, where channel widths measured on average 13.78 *μ*m under glucose limitation and 11.27 *μ*m under ammonium limitation.

**Supplementary Figure 5:**
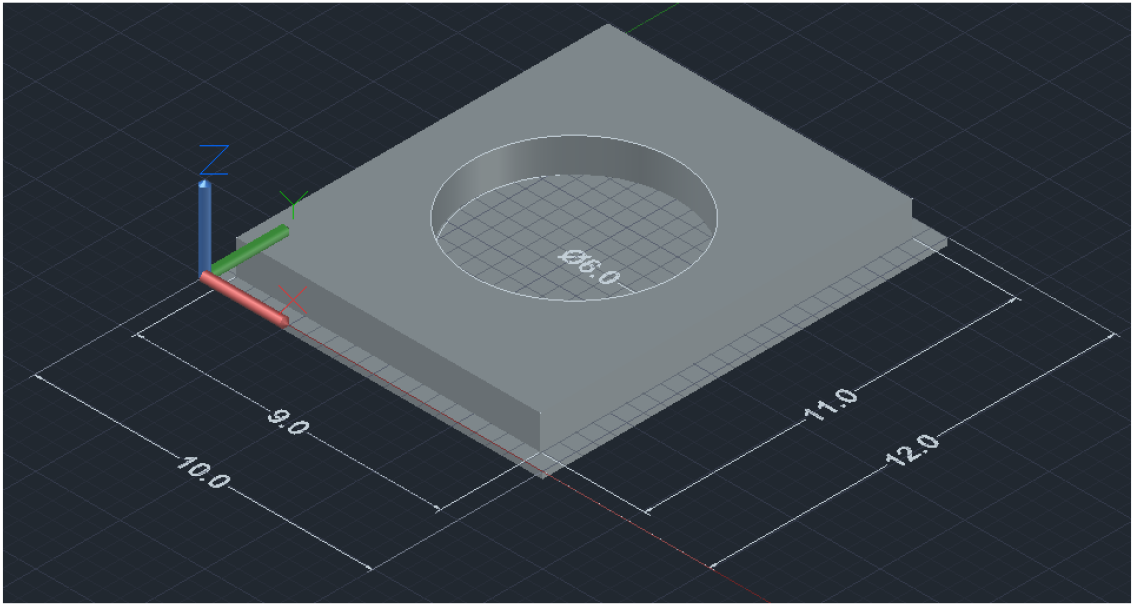
3D-printed imaging chamber used with the Mesolens. Measurements are given in cm.

## References

[1] Donlan RM. 2002 Biofilms: microbial life on surfaces. Emerging infectious diseases 8, 9, 881. (doi:10.3201/eid0809.020063).

[2] Yin W, Wang Y, Liu L, He J. 2019 Biofilms: the microbial “protective clothing” in extreme environments. International journal of molecular sciences 20, 14, 3423. (doi:10.3390/ijms20143423).

[3] Cont A, Rossy T, Al-Mayyah Z, Persat A. 2020 Biofilms deform soft surfaces and disrupt epithelia. Elife 9, e56533. (doi:10.7554/eLife.56533).

[4] Mimura M, Sakaguchi H, Matsushita M. 2000 Reaction–diffusion modelling of bacterial colony patterns. Physica A: Statistical Mechanics and its Applications 282, 1-2, 283–303. (doi:10.1016/S0378-4371(00)00085-6).

[5] Tokita R, Katoh T, Maeda Y, Wakita Ji, Sano M, Matsuyama T, Matsushita M. 2009 Pattern formation of bacterial colonies by escherichia coli. Journal of the Physical Society of Japan 78, 7, 074005–074005. (doi:10.1143/JPSJ.78.074005).

[6] Fei C, Mao S, Yan J, Alert R, Stone HA, Bassler BL, Wingreen NS, Košmrlj A. 2020 Nonuniform growth and surface friction determine bacterial biofilm morphology on soft substrates. Proceedings of the National Academy of Sciences 117, 14, 7622–7632. (doi:10.1073/pnas.1919607117).

[7] Bryers JD. 2008 Medical biofilms. Biotechnology and bioengineering 100, 1, 1–18. (doi:10.1002/bit.21838).

[8] Song F, Ren D. 2014 Stiffness of cross-linked poly (dimethylsiloxane) affects bacterial adhesion and antibiotic susceptibility of attached cells. Langmuir 30, 34, 10354–10362. (doi:10.1021/la502029f).

[9] Kolewe KW, Peyton SR, Schiffman JD. 2015 Fewer bacteria adhere to softer hydrogels. ACS applied materials & interfaces 7, 35, 19562–19569. (doi:10.1021/acsami.5b04269).

[10] Song F, Brasch ME, Wang H, Henderson JH, Sauer K, Ren D. 2017 How bacteria respond to material stiffness during attachment: a role of escherichia coli flagellar motility. ACS applied materials & interfaces 9, 27, 22176–22184. (doi:10.1021/acsami.7b04757).

[11] Saha N, Monge C, Dulong V, Picart C, Glinel K. 2013 Influence of polyelectrolyte film stiffness on bacterial growth. Biomacromolecules 14, 2, 520–528. (doi:10.1021/bm301774a).

[12] Grant MA, Wacław B, Allen RJ, Cicuta P. 2014 The role of mechanical forces in the planar-to-bulk transition in growing escherichia coli microcolonies. Journal of The Royal Society Interface 11, 97, 20140400. (doi:10.1098/rsif.2014.0400).

[13] Paliy O, Gunasekera TS. 2007 Growth of e. coli bl21 in minimal media with different gluconeogenic carbon sources and salt contents. Applied microbiology and biotechnology 73, 5, 1169–1172. (doi:10.1007/s00253-006-0554-8).

[14] Shehata TE, Marr AG. 1971 Effect of nutrient concentration on the growth of escherichia coli. Journal of bacteriology 107, 1, 210–216. (doi:10.1128/jb.107.1.210-216.1971).

[15] Bauchop T, Elsden S. 1960 The growth of micro-organisms in relation to their energy supply. Microbiology 23, 3, 457–469. (doi:10.1099/00221287-23-3-457).

[16] Kandemir N, Vollmer W, Jakubovics NS, Chen J. 2018 Mechanical interactions between bacteria and hydrogels. Scientific reports 8, 1, 1–11. (doi:10.1038/s41598-018-29269-x).

[17] Witten TA, Sander LM. 1983 Diffusion-limited aggregation. Physical review B 27, 9, 5686. (doi:10.1103/PhysRevB.27.5686).

[18] Fujikawa H, Matsushita M. 1991 Bacterial fractal growth in the concentration field of nutrient. Journal of the Physical Society of Japan 60, 1, 88–94. (doi:10.1143/JPSJ.60.88).

[19] Beyenal H, Lewandowski Z. 2000 Combined effect of substrate concentration and flow velocity on effective diffusivity in biofilms. Water research 34, 2, 528–538. (doi:10.1016/s0043-1354(99)00147-5).

[20] Díaz-Pascual F, Lempp M, Nosho K, Jeckel H, Jo JK, Neuhaus K, Hartmann R, Jelli E, Hansen MF, Price-Whelan A, et al. 2021 Spatial alanine metabolism determines local growth dynamics of escherichia coli colonies. Elife 10, e70794. (doi:10.7554/eLife.70794).

[21] Wolfsberg E, Long CP, Antoniewicz MR. 2018 Metabolism in dense microbial colonies: 13c metabolic flux analysis of e. coli grown on agar identifies two distinct cell populations with acetate cross-feeding. Metabolic engineering 49, 242–247. (doi:10.1016/j.ymben.2018.08.013).

[22] Benomar S, Ranava D, Cárdenas ML, Trably E, Rafrafi Y, Ducret A, Hamelin J, Lojou E, Steyer JP, Giudici-Orticoni MT. 2015 Nutritional stress induces exchange of cell material and energetic coupling between bacterial species. Nature communications 6, 1, 1–10. (doi:10.1038/ncomms7283).

[23] Massol-Deyá AA, Whallon J, Hickey RF, Tiedje JM. 1995 Channel structures in aerobic biofilms of fixed-film reactors treating contaminated groundwater. Appl. Environ. Microbiol. 61, 2, 769–777. (doi:10.1128/aem.61.2.769-777.1995).

[24] Tolker-Nielsen T, Molin S. 2000 Spatial organization of microbial biofilm communities. Microbial ecology 40, 2, 75–84. (doi:10.1007/s002480000057).

[25] Birjiniuk A, Billings N, Nance E, Hanes J, Ribbeck K, Doyle PS. 2014 Single particle tracking reveals spatial and dynamic organization of the escherichia coli biofilm matrix. New journal of physics 16, 8, 085014. (doi:10.1088/1367-2630/16/8/085014).

[26] Rooney LM, Amos WB, Hoskisson PA, McConnell G. 2020 Intra-colony channels in e. coli function as a nutrient uptake system. The ISME journal 14, 10, 2461–2473. (doi:10.1038/s41396-020-0700-9).

[27] McConnell G, Trägårdh J, Amor R, Dempster J, Reid E, Amos WB. 2016 A novel optical microscope for imaging large embryos and tissue volumes with sub-cellular resolution throughout. Elife 5, e18659. (doi:10.7554/eLife.18659).

[28] Chen G, Strevett K. 2003 Impact of carbon and nitrogen conditions on e. coli surface thermodynamics. Colloids and Surfaces B: Biointerfaces 28, 2–3, 135–146. (doi:10.1016/S0927-7765(02)00143-1).

[29] Warren MR, Sun H, Yan Y, Cremer J, Li B, Hwa T. 2019 Spatiotemporal establishment of dense bacterial colonies growing on hard agar. Elife 8, e41093. (doi:10.7554/eLife.41093).

[30] Wimpenny JW. 1979 The growth and form of bacterial colonies. Microbiology 114, 2, 483–486. (doi:10.1099/00221287-114-2-483).

[31] Gingichashvili S, Feuerstein O, Steinberg D. 2021 Topography and expansion patterns at the biofilm-agar interface in bacillus subtilis biofilms. Microorganisms 9, 1, 84. (doi:10.3390/microorganisms9010084).

[32] Yan J, Fei C, Mao S, Moreau A, Wingreen NS, Košmrlj A, Stone HA, Bassler BL. 2019 Mechanical instability and interfacial energy drive biofilm morphogenesis. Elife 8, e43920. (doi:10.7554/eLife.43920).

[33] Nijjer J, Li C, Zhang Q, Lu H, Zhang S, Yan J. 2021 Mechanical forces drive a reorientation cascade leading to biofilm self-patterning. Nature communications 12, 1, 1–9. (doi:10.1038/s41467-021-26869-6).

[34] Qin B, Fei C, Bridges AA, Mashruwala AA, Stone HA, Wingreen NS, Bassler BL. 2020 Cell position fates and collective fountain flow in bacterial biofilms revealed by light-sheet microscopy. Science 369, 6499, 71–77. (doi:10.1126/science.abb8501).

[35] Serra DO, Hengge R. 2021 Bacterial multicellularity: The biology of escherichia coli building large-scale biofilm communities. Annual review of microbiology 75, 269–290. (doi:10.1146/annurev-micro-031921-055801).

[36] Yao Z, Davis RM, Kishony R, Kahne D, Ruiz N. 2012 Regulation of cell size in response to nutrient availability by fatty acid biosynthesis in escherichia coli. Proceedings of the National Academy of Sciences 109, 38, E2561–E2568. (doi:10.1073/pnas.1209742109).

[37] Chanson H. 2004 >Hydraulics of open channel flow. Elsevier, 2 ed. (doi:10.1016/B978-075065978-9/50020-9).

[38] Batchelor C, Batchelor G. 2000 An introduction to fluid dynamics. Cambridge university press.

[39] Waldrop R, McLaren A, Calara F, McLemore R. 2014 Biofilm growth has a threshold response to glucose in vitro. Clinical Orthopaedics and Related Research^®^ 472, 11, 3305–3310. (doi:10.1007/s11999-014-3538-5).

[40] Pirt S. 1967 A kinetic study of the mode of growth of surface colonies of bacteria and fungi. Microbiology 47, 2, 181–197. (doi:10.1099/00221287-47-2-181).

[41] Chen L, Noorbakhsh J, Adams RM, Samaniego-Evans J, Agollah G, Nevozhay D, Kuzdzal-Fick J, Mehta P, Balázsi G. 2014 Two-dimensionality of yeast colony expansion accompanied by pattern formation. PLoS computational biology 10, 12, e1003979. (doi:10.1371/journal.pcbi.1003979).

[42] Chacón JM, Möbius W, Harcombe WR. 2018 The spatial and metabolic basis of colony size variation. The ISME journal 12, 3, 669–680. (doi:10.1038/s41396-017-0038-0).

[43] Yang A, Tang WS, Si T, Tang JX. 2017 Influence of physical effects on the swarming motility of pseudomonas aeruginosa. Biophysical Journal 112, 7, 1462–1471.

[44] Reshes G, Vanounou S, Fishov I, Feingold M. 2008 Timing the start of division in e. coli: a single-cell study. Physical biology 5, 4, 046001. (doi:10.1088/1478-3975/5/4/046001).

[45] Kreft JU, Picioreanu C, Wimpenny JW, van Loosdrecht MC. 2001 Individual-based modelling of biofilms. Microbiology 147, 11, 2897–2912. (doi:10.1099/00221287-147-11-2897).

[46] Alpkvist E, Picioreanu C, van Loosdrecht MC, Heyden A. 2006 Three-dimensional biofilm model with individual cells and continuum eps matrix. Biotechnology and bioengineering 94, 5, 961–979. (doi:10.1002/bit.20917).

[47] Lardon LA, Merkey BV, Martins S, Dötsch A, Picioreanu C, Kreft JU, Smets BF. 2011 idynomics: next-generation individual-based modelling of biofilms. Environmental microbiology 13, 9, 2416–2434. (doi:10.1111/j.1462-2920.2011.02414.x).

[48] Xavier JB, Picioreanu C, Van Loosdrecht MC. 2005 A framework for multidimensional modelling of activity and structure of multispecies biofilms. Environmental microbiology 7, 8, 1085–1103. (doi:10.1111/j.1462-2920.2005.00787.x).

[49] Yan J, Sharo AG, Stone HA, Wingreen NS, Bassler BL. 2016 Vibrio cholerae biofilm growth program and architecture revealed by single-cell live imaging. Proceedings of the National Academy of Sciences 113, 36, E5337–E5343. (doi:10.1073/pnas.1611494113).

[50] Hartmann R, Singh PK, Pearce P, Mok R, Song B, Díaz-Pascual F, Dunkel J, Drescher K. 2019 Emergence of threedimensional order and structure in growing biofilms. Nature physics 15, 3, 251–256. (doi:10.1038/s41567-018-0356-9).

[51] Drescher K, Dunkel J, Nadell CD, Van Teeffelen S, Grnja I, Wingreen NS, Stone HA, Bassler BL. 2016 Architectural transitions in vibrio cholerae biofilms at single-cell resolution. Proceedings of the National Academy of Sciences 113, 14, E2066–E2072. (doi:10.1073/pnas.1601702113).

[52] Lambertsen L, Sternberg C, Molin S. 2004 Mini-tn7 transposons for site-specific tagging of bacteria with fluorescent proteins. Environmental microbiology 6, 7, 726–732. (doi:10.1111/j.1462-2920.2004.00605.x).

[53] Elbing KL, Brent R. 2019 Recipes and tools for culture of escherichia coli. Current protocols in molecular biology 125, 1, e83. (doi:10.1002/cpmb.83).

[54] Schniete J, Franssen A, Dempster J, Bushell TJ, Amos WB, McConnell G. 2018 Fast optical sectioning for widefield fluorescence mesoscopy with the mesolens based on hilo microscopy. Scientific reports 8, 1, 1–10. (doi:10.1038/s41598-018-34516-2).

[55] Schindelin J, Arganda-Carreras I, Frise E, Kaynig V, Longair M, Pietzsch T, Preibisch S, Rueden C, Saalfeld S, Schmid B, et al. 2012 Fiji: an open-source platform for biological-image analysis. Nature methods 9, 7, 676–682. (doi:10.1038/nmeth.2019).

[56] Donnelly E, Mothe F. 2007. Polar transformer. https://imagej.nih.gov/ij/plugins/polar-transformer.html.

[57] Iglewicz B, Hoaglin DC. 1993 How to detect and handle outliers, vol. 16. Asq Press.

